# Divergent outcomes of SAK3, a T-type calcium channel enhancer, in two luminal-A type breast cancer cell lines: T-47D and MCF7

**DOI:** 10.1101/2023.11.16.567333

**Authors:** Yashashwini Subbamanda Dinesh, Tharunika Subramanian, Aarushi K Zinzuvadia, Andres D Maturana, Anamika Bhargava

## Abstract

Voltage-gated calcium channels have emerged as promising targets in breast cancer. Breast cancer cell lines are widely used as experimental models. In our pursuit to explore the potential of augmenting T-type voltage-gated calcium channels as a therapeutic approach for breast cancer, we made an unexpected discovery: similar breast cancer subtype cell lines exhibited varying responses to SAK3, a T-type voltage-gated calcium channel enhancer, in terms of proliferation and intracellular calcium levels. In presenting these contrasting findings here, we aim to underscore the importance of exercising caution and validating results obtained from cell lines by cross-referencing them with other cell lines or models.

## Introduction

Breast cancer is primarily categorized into four molecular subtypes: basal-like/triple-negative breast cancer (TNBC), luminal A, luminal B, and human epidermal growth factor receptor two positive (HER2+) [1]. In the context of breast cancer proliferation, voltage-gated T-type calcium channels (TTCCs) have received attention as pivotal components in maintaining calcium balance due to the significant role of calcium in cancer development [2]. The three TTCC isoforms, Ca_V_3.1, Ca_V_3.2, and Ca_V_3.3, are encoded by three genes, *CACNA1G, CACNA1H*, and *CACNA1I*. Ca_V_3.1 acts as a tumor suppressor and is downregulated in luminal-type breast cancer [3,4]. Hence, we hypothesized that enhancing Ca_V_3.1 activity could hold therapeutic promise for breast cancer.

In this study, we used two breast cancer model cell lines, T-47D and MCF7 cell lines, both expressing TTCC isoforms [5]. We examined the impact of SAK3, a TTCC enhancer, on the growth rate and intracellular calcium levels of these two cell lines. SAK3 has been reported to increase specifically Ca_V_3.1 and Ca_V_3.3-elicited calcium currents at concentrations ranging from 0.10 to 10 nM in Neuro-2a cells [6]. Interestingly, despite their shared origin and classification as luminal-A breast cancer subtype, these two cell lines, T-47D and MCF7 exhibited contrasting responses to SAK3. Our results suggest that enhancing the activity of the TTCC isoform Ca_V_3.1 with SAK3 shows promise as a therapeutic approach, but that depends on the cell line used. Therefore, we advise caution when interpreting data from various cell lines across multiple studies.

## Materials and methods

### Cell culture, SAK3 treatment and cell proliferation assay

Human luminal A-type breast cancer cell lines MCF7 and T-47D were obtained from the National Centre for Cell Sciences (NCCS, Pune, India) cell repository and maintained under standard culture conditions as described before [4]. Both cell lines were seeded into 24-well plates and 72 h post treatment with SAK3, cell proliferation was determined either by SRB [7] or MTT [4] assay.

### Quantitative PCR (qPCR) experiments

qPCR was done using QuantStudio 3 Real-Time PCR system (Applied Biosystems), as described in detail in the supplementary information. Table S1 shows the primers used in the study.

### SAK3 treatment and intracellular calcium imaging

T-47D and MCF7 cells were seeded onto poly-D-lysine coated coverslips. 24 h after plating 1 nM SAK3 was added. 24 h post SAK3 treatment, cells were loaded with 5 µM Fura-2AM (Invitrogen) for 30 min. Fluorescent imaging was performed to assess intracellular calcium homeostasis. To understand the acute effects of SAK3 on the intracellular calcium, cells were perfused with 1 nM SAK3. Measurements were done in time-lapse mode using Patch Master ^TM^ software. The ratio of 510 nm emission upon excitation at 340 nm and 380 nm was used to denote intracellular calcium in arbitrary values.

### Statistical analysis

All the statistics were performed using the GraphPad Prism software version 8.0. All data are presented as mean ± SEM. * denotes P ≤ 0.05, ** denotes P ≤ 0.001 and *** denotes P ≤ 0.0001.

## Results and discussion

### 1. qPCR reveals expression of all isoforms of the TTCC gene in T-47D and MCF7 breast cancer cell lines

We observed expression of all three isoforms *CACNA1G, CACNA1H*, and *CACNA1I* in both cell lines, albeit at differential levels (Fig. 1). Ca_V_3.2 has the highest expression, followed by Ca_V_3.3 while Ca_V_3.1 remained at minimal levels in both cell lines. However, we did not observe any significant differences in the expression of TTCC isoforms between the two cell lines. Our results are in agreement with previously published data [8,9] and with data from online patient repositories where Ca_V_3.1 expression is downregulated in breast cancer patients [4] as opposed to other isoforms.

**Figure 1:**
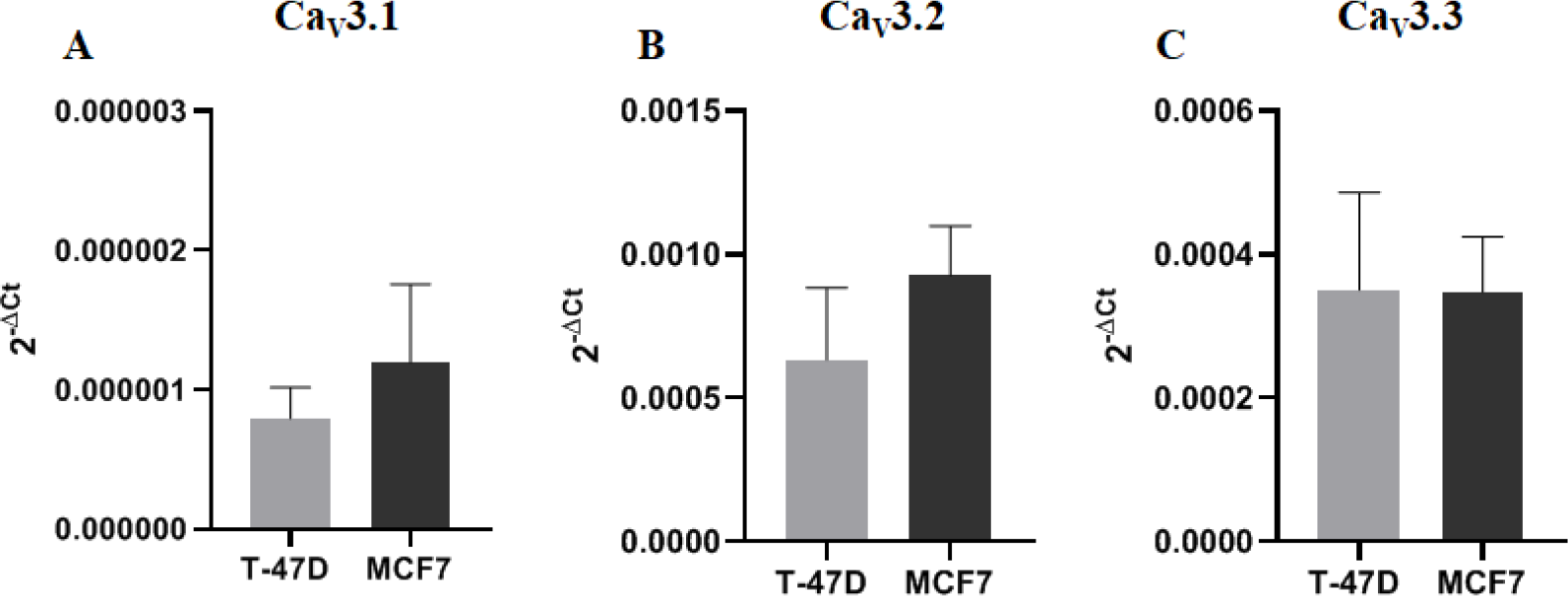
qPCR for TTCC gene isoforms in T-47D and MCF7 cell lines. The expression of T-type calcium channel isoforms in luminal A-type breast cancer cell lines is plotted relative to the β-actin gene. 2^-ΔCt^ was analysed from 4-6 biological replicates.

### 2. SAK3 suppresses the proliferation of MCF7 but not T-47D cells

Treatment of T-47D cells with SAK3 in a concentration range (0.1 to 10 nM) where it would largely enhance Ca_V_3.1, did not produce any significant changes in cell proliferation (Fig. 2A). On the other hand, treatment of MCF7 cells with 1 and 10 nM SAK3 resulted in a significant decrease in cell proliferation (Fig. 2B). This led us to hypothesize that T-47D and MCF7 cells may have disparate calcium signalling upon SAK3 treatment as TTCCs are involved in calcium homeostasis and cell proliferation.

**Figure 2:**
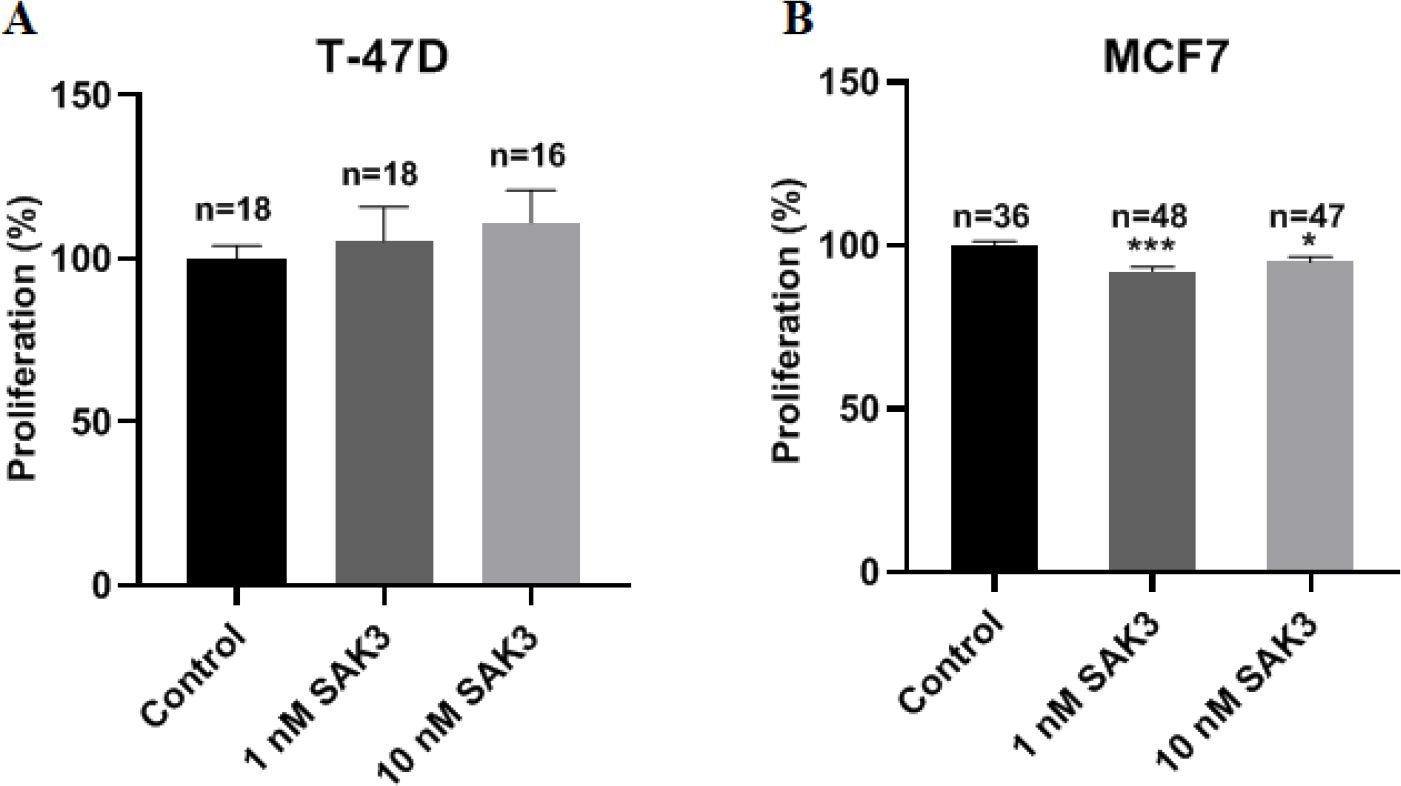
Effect of SAK3 on the proliferation of luminal-A type breast cancer cells. (A) Cellular proliferation after treatment of (A) T-47D or (B) MCF7 cells with SAK3. The ‘n’ numbers above each bar represent the number of wells analysed from three experiments.

### 3. SAK3 treatment led to increased cytosolic calcium in T-47D cells, whereas cytosolic calcium was decreased in MCF7 cells upon SAK3 treatment

To determine whether the differential effect of SAK3 on the proliferation of T-47D and MCF7 cells could be attributed to changes in the intracellular calcium, the cytosolic calcium level was measured in T-47D and MCF7 cells upon treatment with 1 nM SAK3. We used 1 nM as it demonstrated the most significant impact on the proliferation of MCF7 cells (Fig. 2B). SAK3 treatment led to an increase of the basal cytosolic calcium level in T-47D cells (Fig. 3A), whereas it led to a decrease of the basal cytosolic calcium level in MCF7 cells (Fig. 3B). Hence, we propose that the decrease in MCF7 cell proliferation upon SAK3 treatment may be linked to a reduced cytosolic calcium basal level.

**Figure 3:**
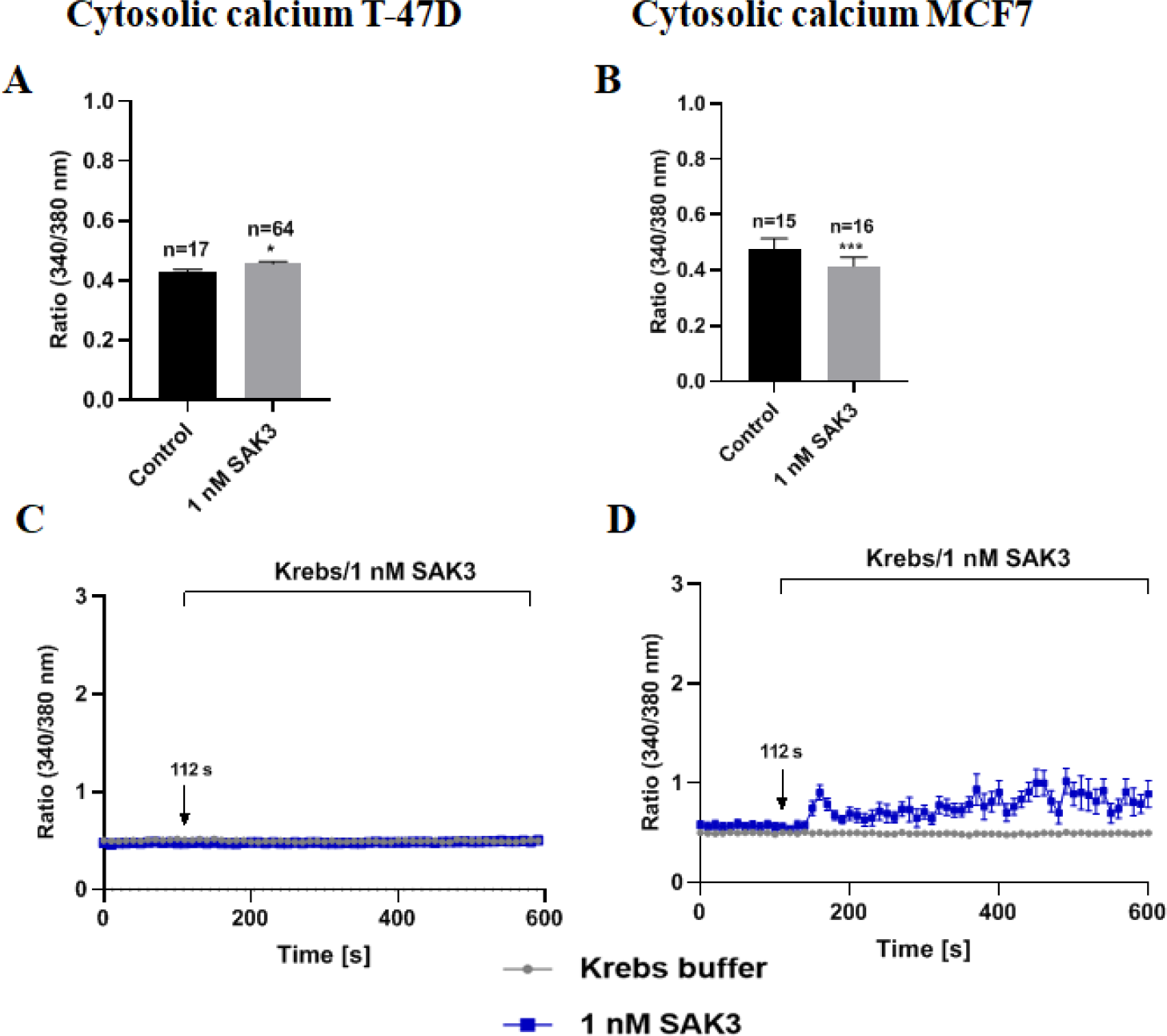
Effect of SAK3 on the cytosolic calcium levels in T-47D and MCF7 cells. Intracellular calcium was measured for 5 s and then averaged as shown in (A) T-47D and (B) MCF7. Intracellular calcium measured during acute application of SAK3 in (C) T-47D or (D) MCF7 cells. The ‘n’ numbers above each bar represent the number of analyzed cells. For acute response (C and D), the number of cells analyzed: Krebs=26, SAK3=43 for T-47D cells, Krebs=9 and SAK3=14 for MCF7 cells.

Since ion channels are transmembrane proteins involved in fast signalling events, we measured the acute response of perfusion of 1 nM SAK3 on the intracellular calcium levels in both cell lines. Perfusion of SAK3 on T-47D cells showed no changes in cytosolic calcium levels, whereas it led to a rapid and notable increase of the cytosolic calcium level in MCF7 cells (Fig. 3C and 3D), indicating that SAK3 modulation of cytosolic calcium levels is more prominent in MCF7 cells.

Overall, our findings reveal that SAK3 influences proliferation and intracellular calcium dynamics differentially in two luminal-A breast cancer cell lines. Further investigations will help to unravel underlying signalling mechanisms and potential therapeutic benefits of SAK3-mediated calcium modulation in breast cancer.

## Supporting information

Supplemental details and Table S1

## Acknowledgements

This work was supported by the IIT Hyderabad, ICMR-DHR Adhoc grants to AB. Japan friendship 2.0 grant to AB and ADM. CSIR, India research fellowship to YSD and AKZ.

## Statements and declarations

### Conflicts of interest/competing interests

The authors declare that they have no conflicts of interest.

### Funding

The authors have no relevant financial or non-financial interests to disclose.

### Ethics approval

Not applicable

### Consent to participate

Not applicable.

### Consent to publish

Not applicable.

### Availability of data and material

All data are available upon request to the corresponding author.

## Author contributions

**YSD**: Experiments, Analysis, Editing, Illustrations, Supervision; **TS:** Experiments, Analysis, Drafting; **AKZ:** Experiments, Analysis, Reviewing; **ADM**: Conceptualization, Reviewing, Funding acquisition, Editing; **AB:** Conceptualization, Analysis, Drafting, Editing, Reviewing, Illustrations, Funding acquisition.

## Notes

### Competing Interest Statement

The authors have declared no competing interest.

