## Supplemental details and Table S1 for "Divergent outcomes of SAK3, a T-type calcium channel enhancer, in two luminal-A type breast cancer cell lines: T-47D and MCF7"

**Supplementary Methods**

1. **cDNA synthesis and quantitative PCR from cell lines**

T-47D and MCF7 cells were maintained according to standard cell culture conditions. Total RNA was isolated from 40-50 % confluence cells using TRIzol reagent following manufacturer’s instructions and quantified by NanoDrop™ 2000 spectrophotometer. The first-strand of cDNA was synthesized by reverse transcription of 1 µg of total RNA using ThermoScientific’s Verso cDNA synthesis kit (AB-1453/B) and was stored at -20 °C till further use. The basal expression of *CACNA1G*, *CACNA1H,* and *CACNA1I* gene was performed on QuantStudio 3 Real-Time PCR system (Applied Biosystems) using Applied Biosystems™ PowerUp^TM^ SYBR^TM^ Green Master Mix (A25742). Primer sequences for the above-mentioned genes are provided in the supplementary table S1. The primers were synthesized by Integrated DNA Technologies (IDT) and supplied by Biosquare Biotechnologies India Private Ltd. The thermal cycling conditions were- UDG activation at 50 °C for 2 min, Dual-Lock DNA polymerase activation at 95 °C for 2 min followed by 45 cycles of denaturation at 95 °C for 15 s, annealing at 55 °C for 15 s and extension at 72 °C for 1 min. Immediately, a default dissociation step was performed to check the non-specific amplification by using the following conditions: 95 °C for 15 s, 60 °C for 1 min and 95 °C for 15 s. In this dissociation step, the temperature was increased gradually and the fluorescence intensity was monitored to generate the dissociation-curve. The mRNA expression levels of above-mentioned genes were normalized to *β-actin* and the relative expression was plotted as 2^-ΔCt^. Each sample was run in triplicates and no-template reactions were used as experimental control to identify PCR contamination.

**Table S1:** Primer sequences of the genes used for qPCR

| **Gene** | **Sequence (5’-3’)** |
| --- | --- |
| ***β-actin*** | FP: CTGCCCTGAGGCACTCTTC  RP: CGGATGTCCACGTCACACTT |
| ***CACNA1G*** | FP: ACCAAGCAGCGGGAAAG  RP: GATGTACACCAGGTACTTGAGC |
| ***CACNA1H*** | FP: GACGCCTTCATTTTCGCCTT  RP: CCCAGGTAACACTTCTGCCC |
| ***CACNA1I*** | FP: GCCCTACTATGCCACCTATTG  RP: AGGCAGATGATGAAGGTGATG |
